# Low plastic ingestion rate in Atlantic Cod (*Gadus morhua*) from Newfoundland destined for human consumption collected through citizen science methods

**DOI:** 10.1101/080986

**Authors:** Max Liboiron, France Liboiron, Emily Wells, Natalie Richárd, Alexander Zahara, Charles Mather, Hillary Bradshaw, Judyannet Murichi

## Abstract

Marine microplastics are a contaminant of concern because their small size allows ingestion by a wide range of marine life. Using citizen science during the Newfoundland recreational cod fishery, we sampled 205 Atlantic cod (*Gadus morhua*) destined for human consumption and found that 5 had eaten plastic, an ingestion prevalence rate of 2.4%. This ingestion rate for Atlantic cod is the second lowest recorded rate in the reviewed published literature (the lowest is 1.4%), and the lowest for any fish in the North Atlantic. This is the first report for plastic ingestion in fish in Newfoundland, Canada, a province dependent on fish for sustenance and livelihoods.

**Highlights (3–5 points, 85 char max including spaces):**

- Plastic ingestion rate of 2.4% for Atlantic Cod (n=205)
- First recorded baseline for fish in Newfoundland, Canada
- This plastic ingestion prevalence rate is among the lowest recorded to date
- Used citizen science to collect GI tracts from fish destined for human consumption

## Introduction

There are plastics in every ocean around the world (Provencher et al. 2010, Zarfl & Matthies 2010, Eriksen et al. 2014) and more than 90% of marine plastics are microplastics (<5mm) (Eriksen et al. 2014). Microplastics (<5mm) and mesoplastics (5–10 mm) are of concern because they are viable for ingestion by a wide range of marine life (Foekema et al. 2013, Wright et al. 2013, Setälä et al. 2014, Kühn et al. 2015). Plastics have been found in marine species that are commonly eaten by humans, including: shrimp (Setälä et al. 2014); bivalves, such as oysters and mussels (Von Moos et al. 2012, Van Cauwenberghe and Janssen 2014); and fish from a variety of trophic levels (Choy and Drazen 2013, Foekema et al. 2013, Jantz et al. 2013, Lusher et al. 2015a, Seltenrich 2015, Bråte et al. 2016). The majority of these studies assess marine life that were raised in a laboratory or caught specifically for research purposes. Few studies have targeted marine life that was caught specifically for human consumption (Van Cauwenberghe and Janssen 2014, Rochman et al. 2015a), and to our knowledge, none sample fish that were eaten by humans.

The study of plastics in food webs is an emerging area of study (Rochman 2016). Research indicates that chemicals accumulate on plastics (Ogata et al. 2009, Mato et al. 2001, Holmes et al. 2012), such as hexachlorinated hexanes (HCHs), polycyclic aromatic hydrocarbons (PAHs), polychlorinated biphenyls (PCBs), polybrominated diphenyl ethers (PBDEs), triclosan, nonylphenols, and heavy metals, among others. The chemicals may then transfer to the animal’s tissues after ingestion (Endo et al. 2005, Browne et al. 2013, Gassel et al. 2013, Tanaka et al. 2013, Rochman et al. 2014, Wardrop et al. 2016). The toxic effects of these synthetic chemicals are varied. For example, they may cause cellular necrosis and tissue lacerations in the gastrointestinal tract (Rochman et al. 2015b), make animals more susceptible to stress (Browne et al. 2013, Rochman et al. 2013), and can result in liver toxicity and pathology (Rochman et al. 2013). While these studies on the harm caused by plastic ingestion are conducted in a laboratory rather than the field, and laboratory contexts reduce the complexity of chemical exposures by studying a single chemical at specific endpoints rather than the “cocktail” of chemicals and wide range of endpoints they may effect (Koelmans et al. 2013, Koelmans et al. 2014), they do provide cause for concern because of the potential human health effects for people who eat fish.

The accumulation of microplastics and their associated toxicants within marine food webs is of special concern in Newfoundland, Canada, where people rely extensively on marine life for food sustenance. In Newfoundland, Atlantic cod (*Gadus morhua*) is an important species to consider because of its cultural and culinary significance. Up to 82% of households along the west coast of Newfoundland report consuming local seafood more than once a week, of which cod is the preferred food choice (Lowitt 2013). Yet, there are no previous studies on plastics ingested by food fish in the region. While commercial harvesting of Atlantic cod is strictly controlled (Bavington 2011) people in Newfoundland and Labrador are able to fish for Atlantic cod during the seasonal recreational cod fishery. Individuals are permitted to catch five fish per person per day, with a maximum of fifteen fish per boat outing (Fisheries and Oceans Canada 2015).

In 2015, the Newfoundland food fishery took place during two one-week periods in the summer and early fall (Schrank and Roy 2013; Fisheries and Oceans Canada 2015). During the September 2015 food fishery, we were present on public fishing wharves near St. John’s, the only major city in the province and thus the area with the greatest population, to obtain the gastrointestinal (GI) tracts from citizen scientists (both commercial and recreational fish harvesters) to monitor the rate of plastic ingestion in fish destined for human consumption. This study joins an emerging trend in microplastic pollution research that evaluates ingestion rates by marine life that is destined for human consumption. All fish were Atlantic cod (*Gadus morhua*). These fish were analyzed in the lab according to standardized methods (van Franeker et al., 2011). Data were used to determine the rate of ingestion between recreational and commercially caught fish, geographical locations where fish had been caught, and types and characteristics of ingested plastics. While there are other dangers associated with marine plastics to fishing communities, such as entanglement and ghost fishing reducing available fish stock (Hall 2000), the purview of this study is on plastic ingestion in fish caught for human consumption.

To contextualize this study, we conducted a literature review of nearly 100 previous studies of marine plastic ingestion by fish from around the world. The following table illustrates published ingestion rates of various fish species using an array of methods. The majority of the studies assessed plastic ingestion rates regionally and as a result the samples included multiple species of fish. For the table we disaggregate the published data and sort the results by fish species where possible. This has resulted in some species appearing to have very small sample sizes, which is an artefact of our disaggregation. We do not provide details on sampling protocols, which may be found in the original cited articles. Species with an ingestion rate of 0% have been excluded from the table.

The 97 reviewed publications on ingestion rates in various fish species in a wide range of geographical locations used different methodologies to assess ingestion rates, which makes the rates not directly comparable, but they do serve as a base to understand and compare ingestion rates in general terms. Studies found that between 0.7–100% (mean 31%) of individual fish within a species had ingested plastics. Of these, Atlantic cod were found to ingest at a rate between 1.4-13% (n=3, mean 5.8%) (Foekema et al 2013, Rummel et al 2015, Brate et al. 2016), and fish species in northern waters fed directly by the Arctic (such as the North Sea) had ingestion rates of <1–13.2% (n=9, mean 5.3%) (Foekema et al 2013, Rummet et al 2015). We expected, then, that we would find a relatively low rate of plastic ingestion in our cod from northern waters (likely less than the mean of 31% for all fish, and closer to the 5.3–6% rate for fish in northern waters or 1.4–15% rate for Atlantic cod).

## Methods

### Collection of Samples

We collected cod fish gastrointestinal (GI) tracts from local fish harvesters as they gutted their fish during the fall food fishery (September 19th, 20th, and 27th, 2015). The provincial Department of Fisheries requires that all fish caught are filleted on land (DFO 2015), making wharves an ideal location for gathering samples. Field station sites were located on wharves with high fishing activity on the eastern coast of Newfoundland, Canada at Petty Harbour and St.Phillip’s Harbour, both of which are withing an hour’s drive of the privince’s capital, St. John’s, the area of highest population in the province.

Both team members and local fish harvesters were well acquainted with what Atlantic cod look like, and all fish sampled were Atlantic cod. We solicited fish GI tracts from local people gutting their fish on public wharves as well as from a commercial fishery in a private dock in Petty Harbour, where we also collected information about the location fish were harvested. We then extracted the GI tract from the esophagus to the anus after fishermen had filleted their fish, and bagged the GI tract contents for laboratory analysis. Each sample was given a unique identifier which was recorded on a master data sheet and on a slip of paper in the sample bag; the identifier consisted of an abbreviation of the location the fish was caught and a number respective the order in which they had been caught as well as the date. For example, the 8th fish caught in Petty Harbour on October 11th, 2015: PH8, Oct. 11/15. GI tracts that were cut open or nicked during the filleting process were discarded. Because the project was designed around public engagement, each fish harvester was asked to provide a name for their donated fish so they could easily locate the results of the data for their specific catch on an online database of results (CLEAR 2015). After bagging the fish intestines, they were placed into a cooler and taken to a laboratory at Memorial University where they were placed in a freezer until further examination. In total, 205 GI tracts were gathered, 188 from the recreational food fishery and 17 from one commercial vessel.

### Laboratory Procedures

Methods follow and adapted standardized protocol for biomonitoring of microplastics in animal GI tracts developed by van Franeker et al. (2011) for birds. We choose this method over others that use KOH (such as Rochman et al. 2015) because, as a laboratory that works with and within local communities, we opt for the most robust methods that use as little specialized chemicals and equipment as possible so citizen scientists can compare their findings to our results. We thawed the GI tracts in cold water for approximately 2 hours prior to dissection. We used a double sieve method, stacking a 4.75mm (#4) mesh stainless steel sieve above a 1mm (#18) mesh stainless steel sieve. These sieves were selected as 1mm is considered the cut off point of ‘large’ microplastics (1.0–5.0mm) (Wagner et al. 2014), and is the size that has been suggested for cod ingestion studies (European Commission 2014). Large microplastics of a 1.0–5.0 mm size range are considered to be the lower limit of what the naked eye and microscope are able to reliably detect, without the use of a spectrometer (Song et al. 2015). The GI tract was placed in the top 5mm sieve and was cut along the stomach and intestines to the anus using fine scissors. We used a wash bottle to gently rinse out the contents into the sieve to remove all mucus and food. Tissues were closely examined for embedded microplastics. We then visually sorted and separated all plastics; the GI tract was placed aside, and we examined the contents of the sieves; microplastics and other anthropogenic materials were removed with tweezers, rinsed, and placed in a Petri dish for closer examination under a microscope. Ingested animals that were intact enough for dissection were given a sub code and then processed separately using the same method as above.

Suspected anthropogenic debris was examined under a dissecting microscope with both reflected oblique and transmitted light (Olympus SZ61, model SZ2-ILST, with a magnification range of 0.5–12x), where we visually sorted microplastics from organic and other anthropogenic debris based on colour, absence or presence of cellular structure, erosion characteristics of plastics (Cocharine et al. 2009), and, if necessary, breaking the objects open to inspect internal structures after they were weighed and measured. All items requiring further inspection for accurate identification were examined under a compound microscope (magnification 10x, 40x). Samples were placed into folded filter paper to dry for a minimum two days or until the weight had stabilized. Once dry, we transferred the plastics, over a Pyrex dish, into pre-labeled scintillation jars.

All plastics recovered were above the size threshold for reliable visual identification (Song et al. 2015) and where simple to identify as plastics with the naked eye and compound microscope (see image below), and secondary methods such as hot wire tests, and Fourier Transform Infrared Spectroscopy (FTIR) analysis, which are usually reserved for fibres or plastics smaller than 1mm, were not necessary. While this is a limitation of the methodology because we do not gain data on the type of polymers recovered, it is also a strength for our region--by using methods that citizen scientists can also employ, we facilitate comparability between future studies in the area. Moreover, a lack of fibres found reduces issues of air contamination of samples, since none of the recovered samples were light enough for air deposition. As such, a standardized protocol (van Franeker et al. 2011) with reliably consident low resolution data and low threat of contamination is more valuable than a variety of studies with varying degrees of resolution that cannot be compared. As there is no baseline data for Newfoundland and Labrador and there is a push in the province for citizen science in the area, this is important methodologically (Liboiron, 2016).

Prevalence rate is reported as the proportion of sampled cod found to have ingested plastic, and arithmetic means (+/−SE) for number of ingested plastic and mass were reported. Following van Franeker (2004), individual items were categorized as industrial plastic (small symmetrical virgin plastic pellets, or ‘nurdles’) or user plastic (categories of: fragments, sheets, threads, foam). Though van Franeker et al. (2011) does not include fibres in their categories, we included them in our analysis, though none were found. The dimensions of each piece were measured using digital callipers (accurate to 0.01 mm). We documented the opacity (transparent or opaque- the ability to see light through the sample when backlit by the microscope), color, and type of erosion. Characteristics of erosion are based on Cochrane et al. 2009, a study of degradation and erosion patterns on beached marine plastics (categories: pitting, adhering particles, grooves, linear fractures, irregular surface). Opacity was determined based on our ability to see through the plastic from light transmitted from the microscope. If there was no light, we considered the plastic to be opaque. Opacity and color were analyzed in the event that cod are attracted to specific visual characteristics, and type and degree of erosion were analyzed to aid in determining whether plastics were potentially local or from other landmasses. Air dried plastics were measured in terms of count and mass (in grams) using a Sartorius electronic weighing scale (accurate to 0.0001 g).

We undertook precautions to avoid cross-contamination. All tools were rinsed or wiped down with water and paper cloths, including the microscope lens and plate, Petri dishes, and sieves. Hands were washed, lab coats worn, and hair was tied back. After each dissection, we closely examined our hands and tools for any microplastics that may have adhered.

## Results

Of the 205 of cod collected, 5 had ingested 7 pieces of plastic between them (range 0 – 2 plastics/fish), an ingestion prevalence rate of 2.4%. However, because our collection method involved local fishermen and women, 27 samples contained only part of the GI tract (usually just the stomach). Because other studies have found that some plastics are excreted by animals that ingest them (Ryan 2015), the protocol we used called for investigating the entire GI tract of fish. If we omit the samples that were missing the lower GI tract from our analysis, we had 177 of fish, 4 of which ingested plastics, an ingestion prevalence rate of 2.3%. Omitting fish whose entire GI tract was not sampled led to a very small underestimation of plastics and a slightly lower rate of prevalence. Given the small difference in results, we have decided to include fish with only part of the GI tract, a 2.4% baseline ingestion rate.

**Table 2:** Description of ingested plastics.

| Fish ID | Type | Mass (g) | Size (longest side) mm | Opacity | Pigmentation | Erosion pattern |
| --- | --- | --- | --- | --- | --- | --- |
| PH67 | Fragment | 0.001 | 4.2 | Opaque | White | pitting, particles, grooves, linear fractures |
| Ph67 | Fragment | 0.0002 | 2 | Opaque | White | pitting, particles, grooves, linear fractures |
| PH92 | Fragment | 0.0018 | 2.8 | Opaque | White | grooves, linear fractures |
| SP141 | Thread | 0.0028 | 9 | Opaque | Green | irregular surface |
| SP31 | Sheet | 0.0017 | 6.5 | Opaque | White | none |
| SP31 | Sheet | 0.0011 | 9.7 | Opaque | White | none |
| SP9 | Thread |  | 8.2 | Opaque | Green | none |

Plastics found in the stomachs were of a variety of types: 2 were film/sheet plastic, 2 were threads, and 3 were fragments. None were industrial pellets (“nurdles”) or fibres. Mean length, width, and height were 6.06 ± 1.17 mm, 0.772 ± 0.398 mm, and 2.23 ± 0.594 mm, respectively (range in the longest dimension was 9.7 mm - 2 mm and the shortest dimension 0.001 mm - 2.5 mm). All samples are above 1mm in size, under which human vision is no longer reliable for identification of plastics, even with the aid of a microscope (Song et al 2015). The mean (+/−SE) mass of ingested plastic, which was based on 6 rather that 7 samples because one sample was lost after size measurement but before weight measurement, was 0.00143 ± 0.00036 g per fish (range 0.0002 g - 0.0028 g). Three of the samples were completely unweathered, while four of the seven (including all fragments) were weathered and showed pitting, grooves, and irregular surfaces (n=4). All were opaque. The two threads were green (the same colour as fishing nets and lines common in the area) and the remaining five items were white.

**Figure 1:**
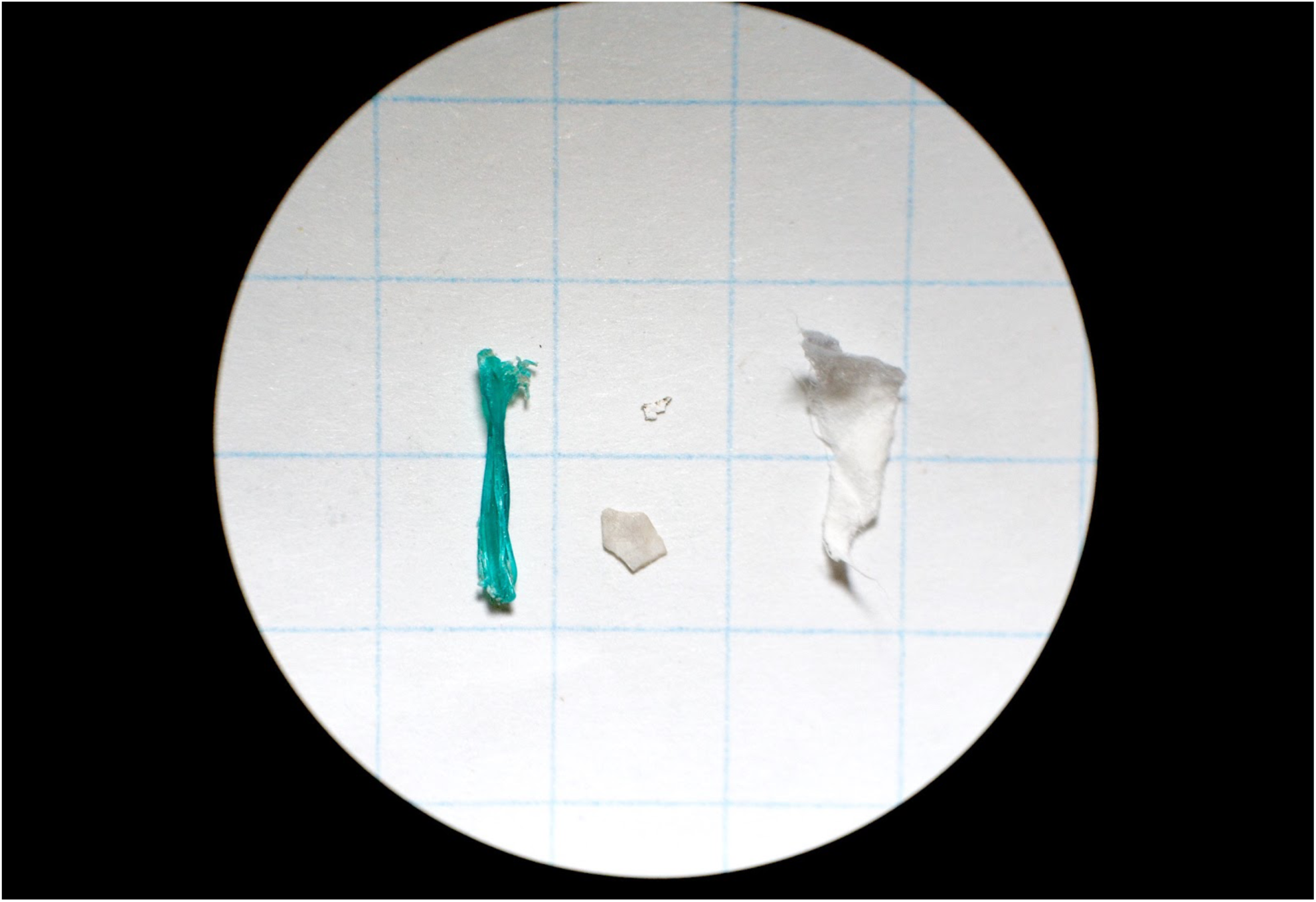
Examples of four plastics found in four Atlantic Cod caught during the Newfoundland recreational cod fishery. The green thread is a typical colour and type found in local fishing gear. The thread (far left) and film (far right) show very little erosion or discolouration, indicating that they have not been in the water long and are likely local plastics. Grid is 1cm x 1cm.

Fish that had ingested plastics came from a variety of locations within an hour’s drive to the capital city of Newfoundland, St. John’s. Two were from Petty Harbour, three from Portugal Cove, one from Quidi Vidi, and one from Bell Island. This distribution is wide given that the vast majority of fish sampled were from Petty Harbour (32.2%) and Portugal Cove (40.5%), with only 11 (5.4%) fish from Belle Island and 3 (1.5%) from Quidi Vidi. Other locations account for the remaining 20.4% of catch. As all samples were collected on wharves at Petty Harbour and St. Phillips, the higher prevalence in samples are from those locations.

**Figure 2:**
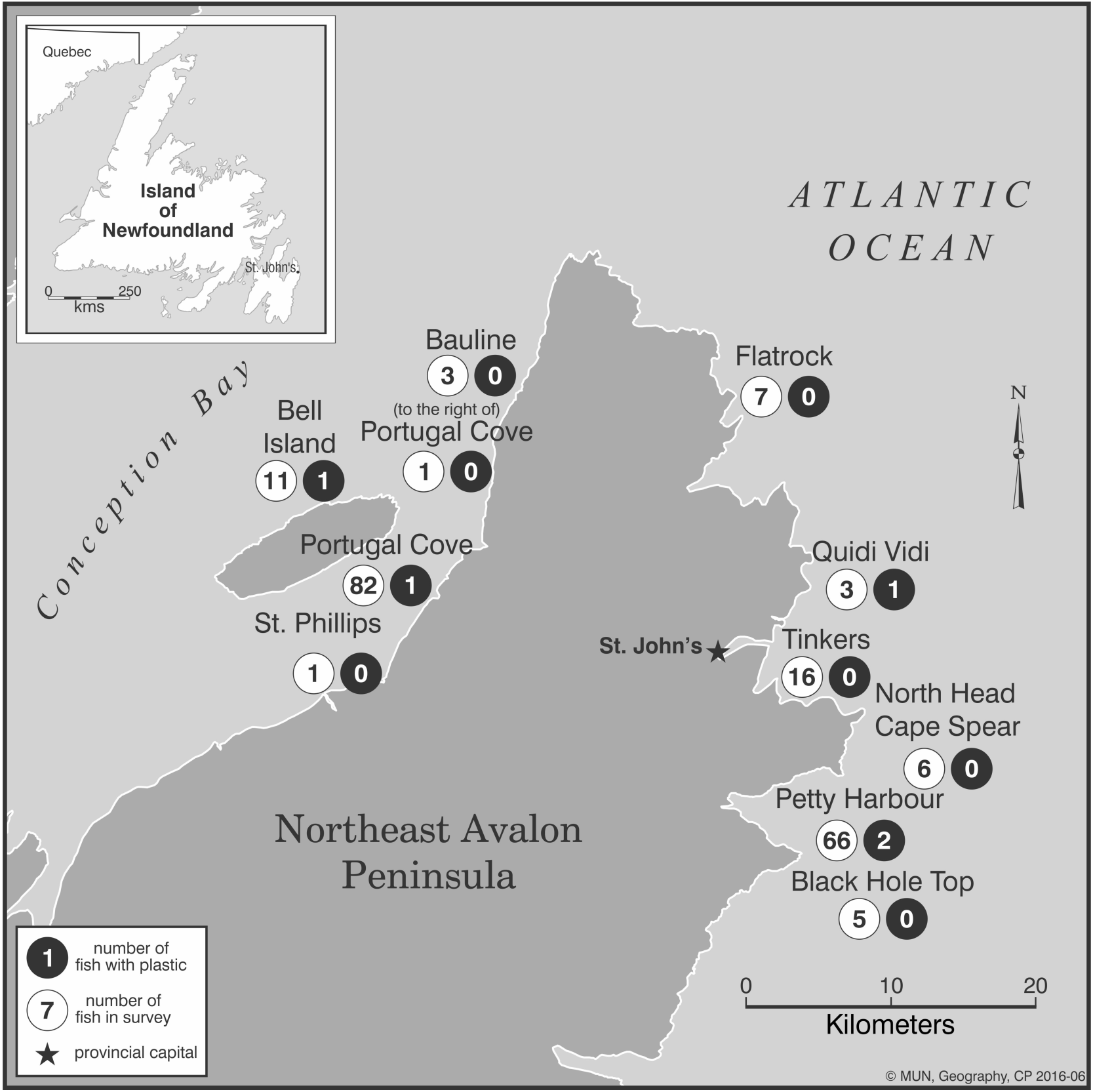
Map of areas in the Avalon Peninsula of Newfoundland where cod fish GI tracts were collected. The number in white indicates the quantity of fish collected from the area, and the number in black indicates how many of those fish had ingested plastic, regardless of the amount of plastics ingested. All fish GI tracts were recovered from the wharves of Petty Harbour or St. Phillips through citizen science methods. All sites are within an hour drive of the province’s capital city, St. John’s.

Most fish sampled (n=188) were from the recreational cod fishery, while 17 were from the commercial fishery, all of which were destined for human consumption. All plastics were found in fish from the recreational fishery. The recreational fishery uses rods and lines to catch fish and stays close to shore (Protected Areas Association of Newfoundland and Labrador 1996). The commercial fishery, which contained no fish that ingested plastic, used bottom trawls further offshore (this is not true of all commercial fisheries in Petty Harbour, but was the case for the fishery we received samples from). All fish that had ingested plastics also had organic food in their stomachs, suggesting a plastic gut clearance rates similar to ingested food (see Brate et al. 2016 for similar findings in Atlantic cod in Norwegian waters).

While the methods for studying ingestion rates in the extensive literature review we conducted were variable, and so direct comparison is not possible, it does help us situate our study compared to other locations and species. Our ingestion rate for Atlantic cod of 2.4% is the second lowest recorded rate in the reviewed literature for cod, below average for fish species in general and fish in northern waters, the lowest for any fish in the North Atlantic.

## Discussion

Since this is the first indication of plastic ingestion rates for a remote province dependant on fish for sustenance and commercial enterprises, we will begin with a discussion of why the ingestion rate may be so low. We hypothesize that Atlantic cod caught for food in Newfoundland have amongst the lowest recorded plastic ingestion rates (2.4%, Table 1) due to three major factors.

**Table 1:**
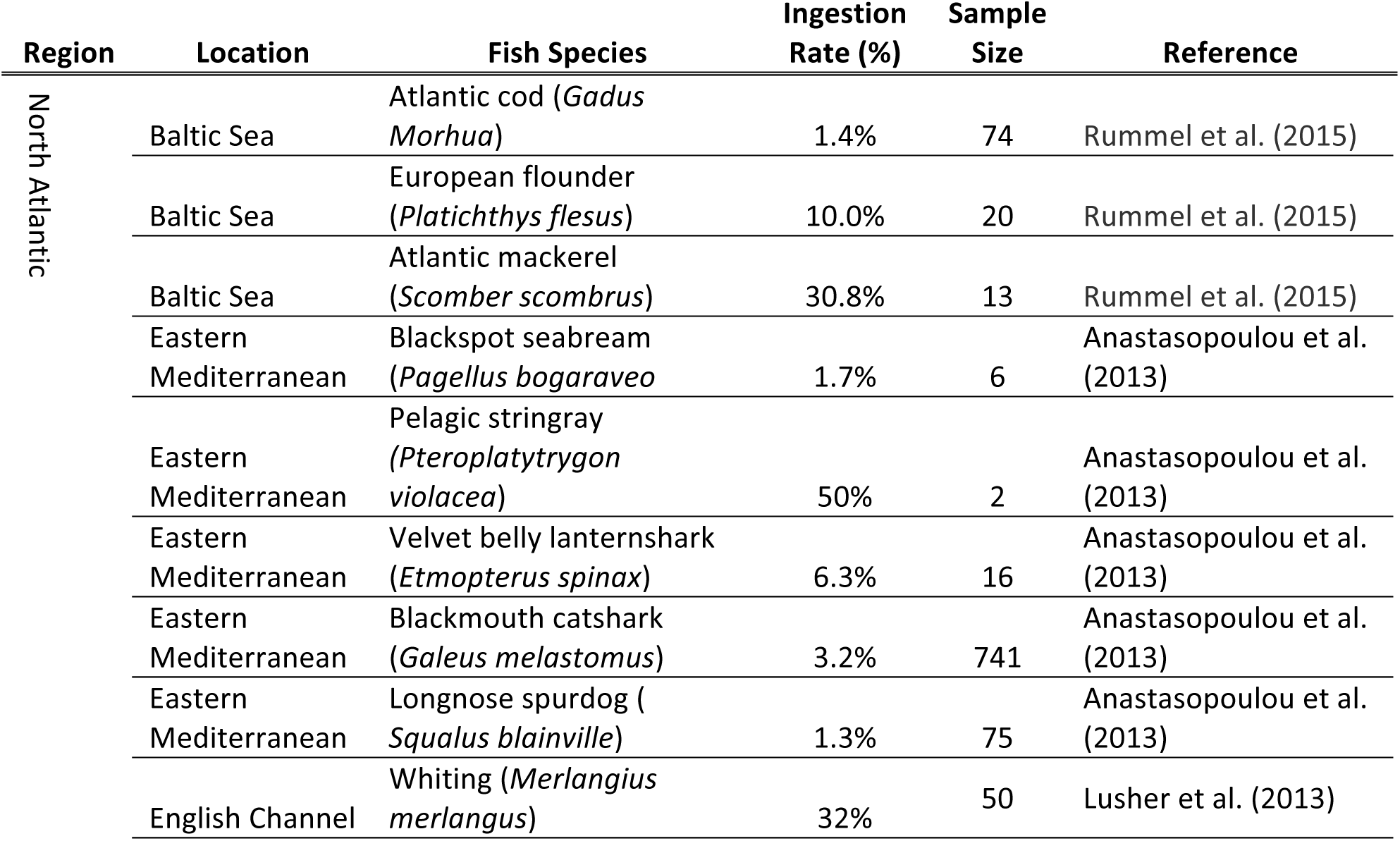
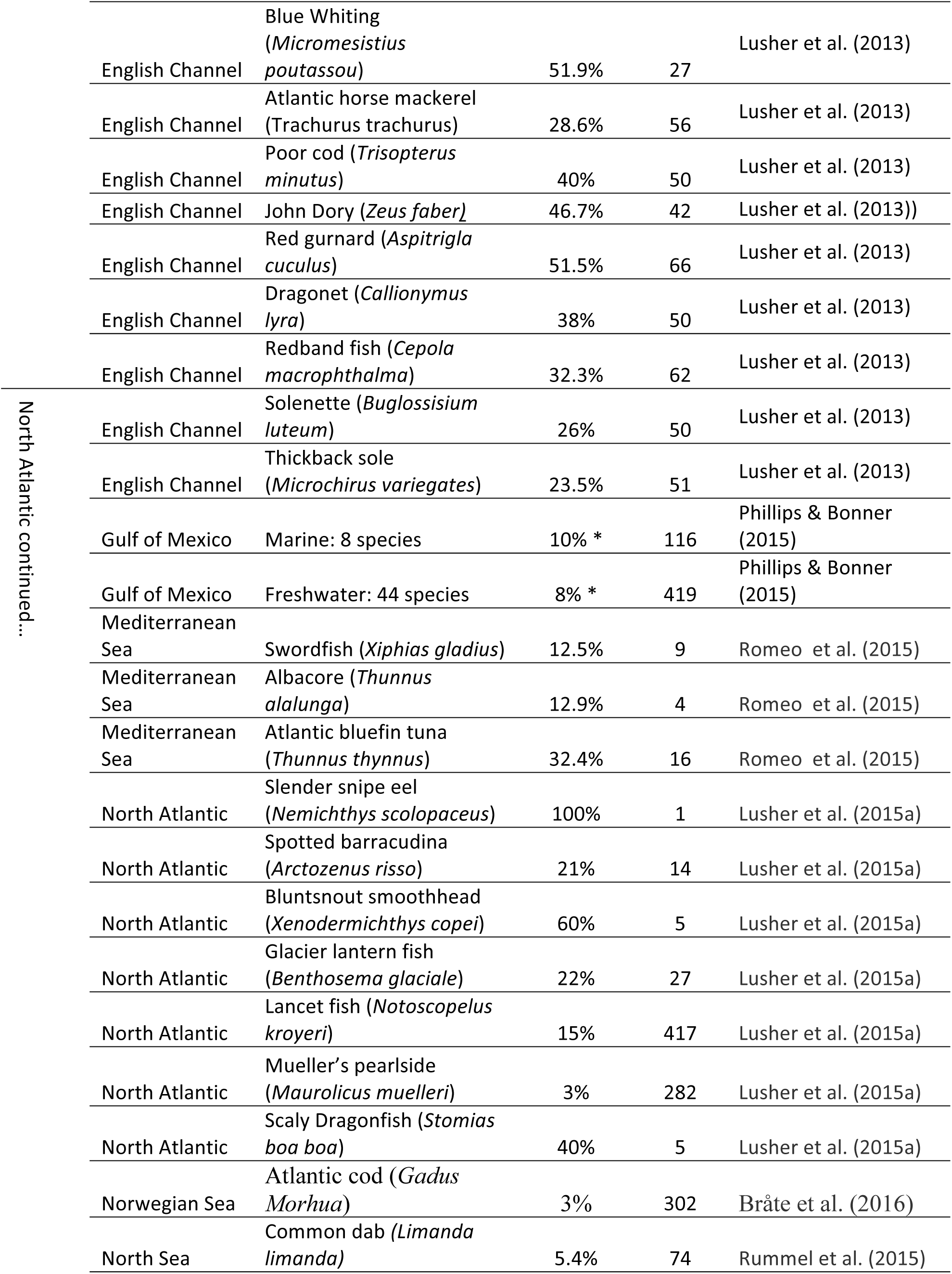
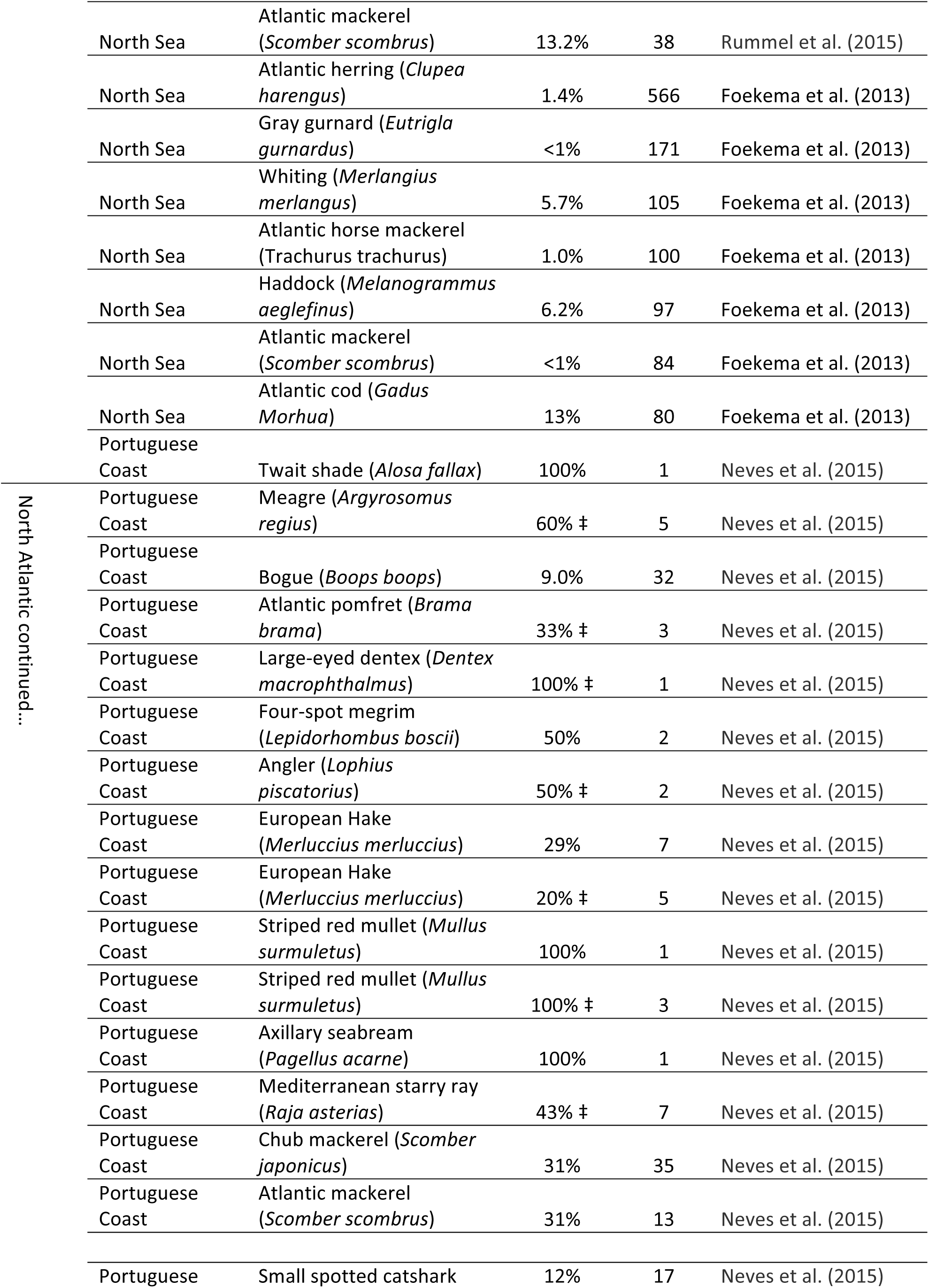
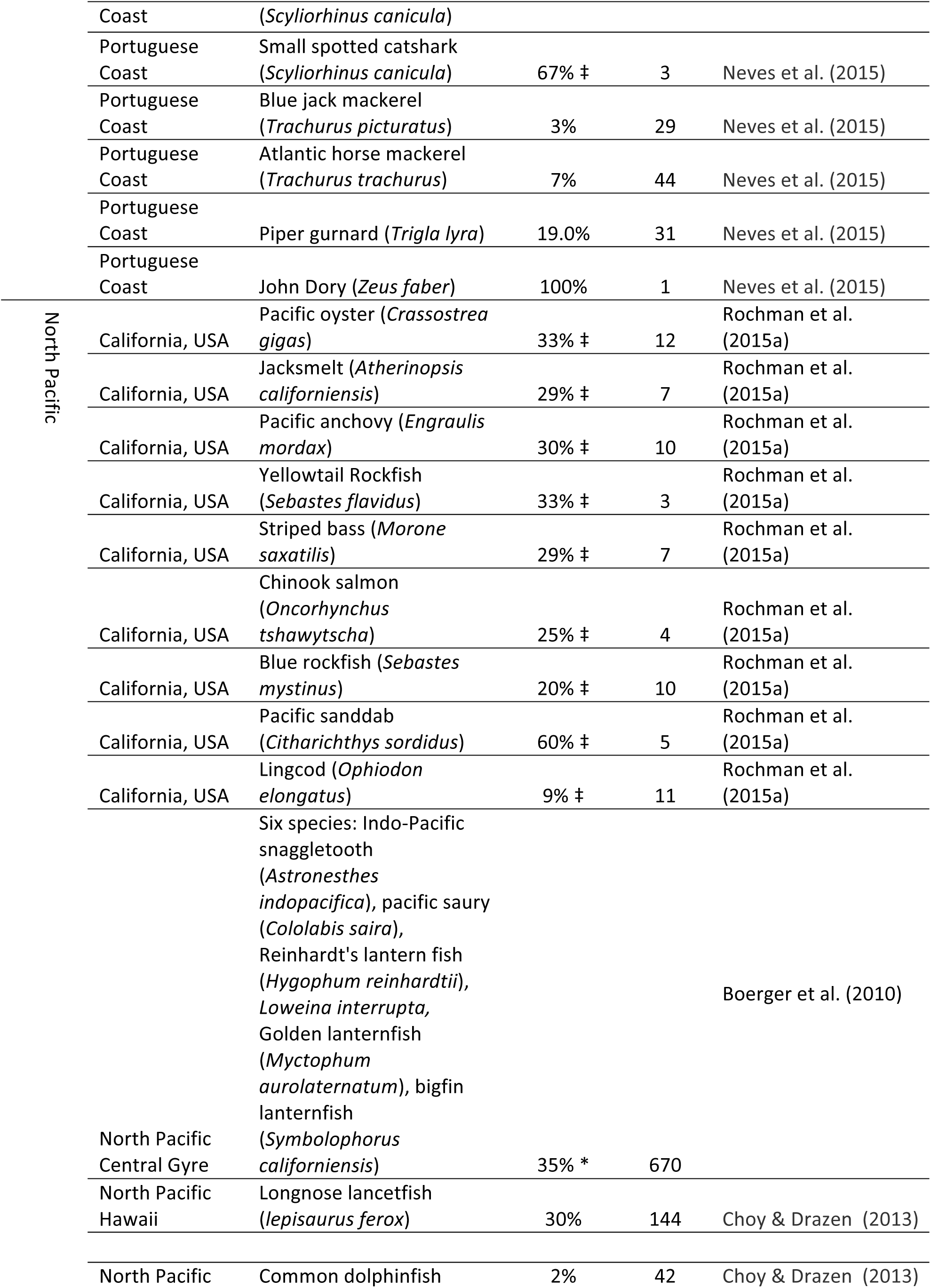
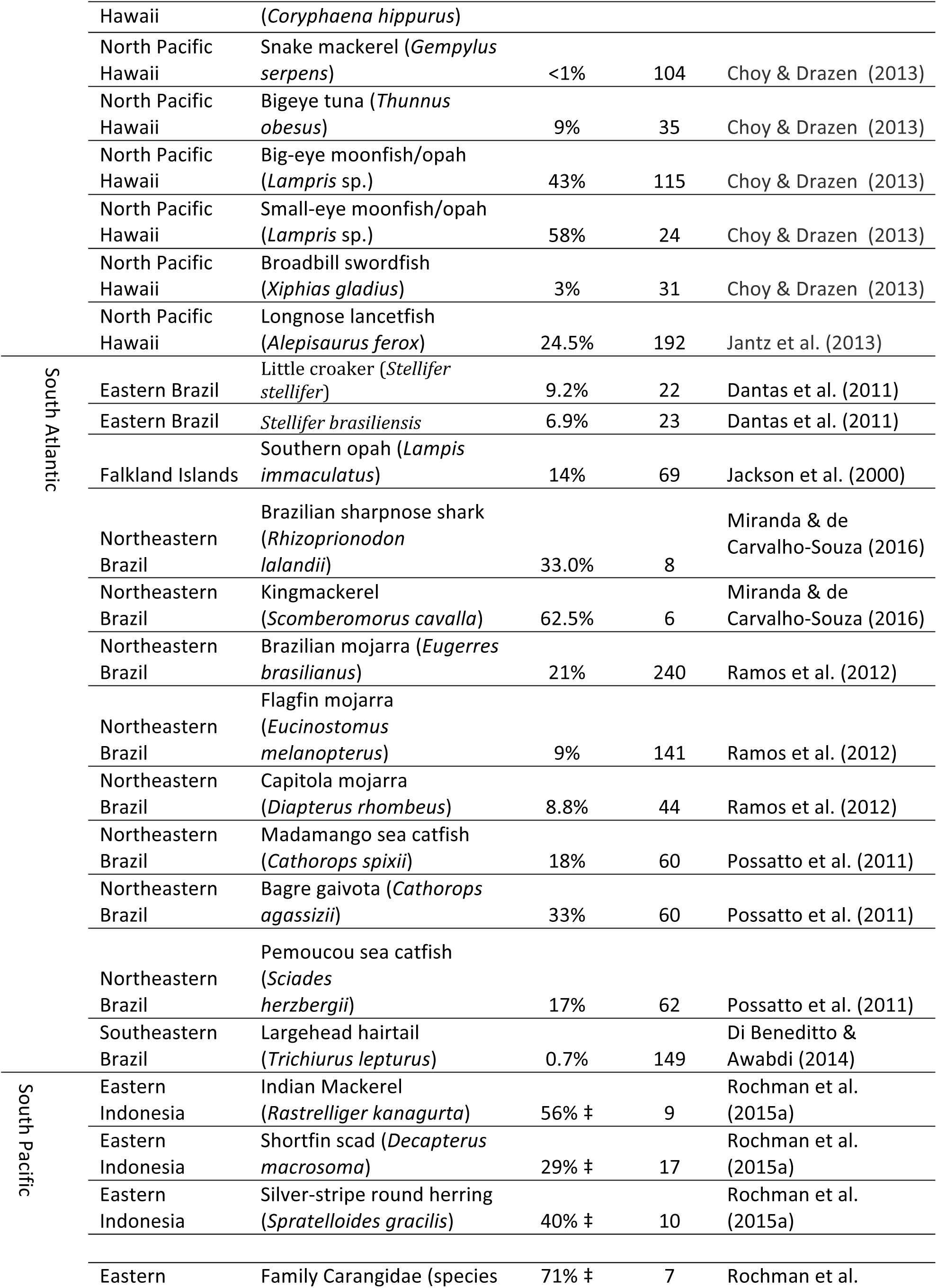
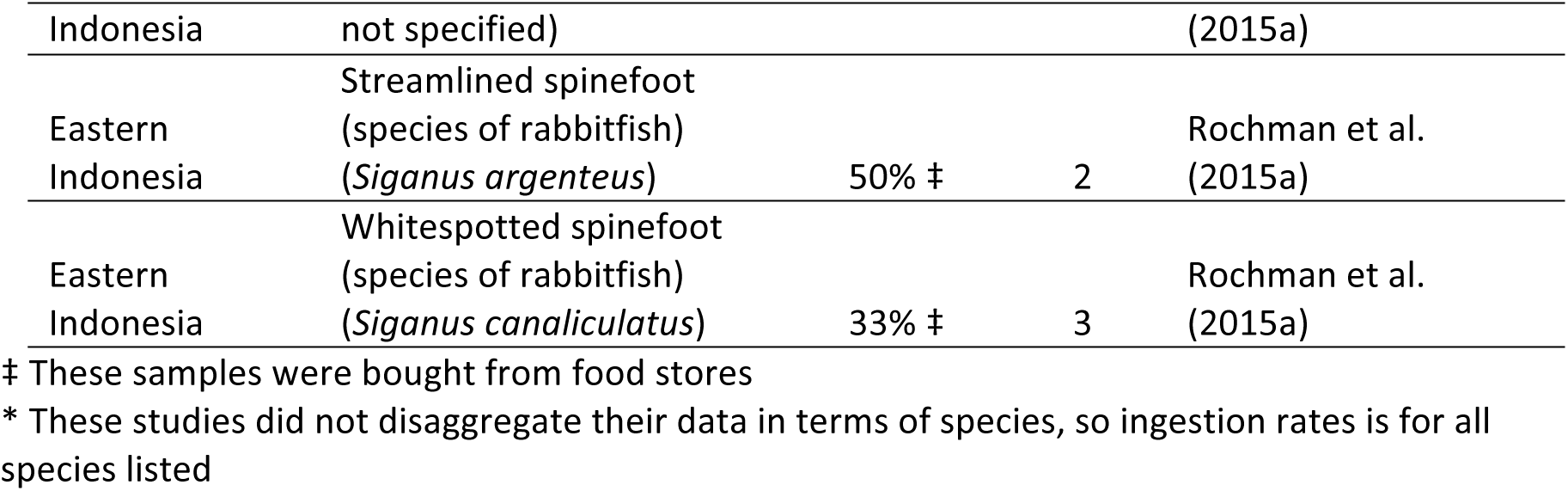
Published plastic ingestion rates of fish, disaggregated by species and body of water.

These factors are: (1) fish consumption behaviour; (2) the prevalence of ocean plastics in Newfoundland waters; (3) the relatively small population of the island of Newfoundland. We then discuss these hypotheses to characterise potential sources of, and susceptibility to, plastic pollution, and to point to future directions for marine plastic research in Newfoundland.

First, we speculate that cod behaviour and feeding patterns make them less likely to encounter ocean plastics than other fish. Understanding ingestion rates within the context of a species life history is important given calls by marine plastics researchers to consider animal behaviour within their study design and analysis (Carson 2013). While research has not determined the abundance of microplastics throughout the water column in the North Atlantic, characteristics of microplastics correspond with their abundance in certain locations: namely, low-density plastics are commonly found near surface or subsurface waters (Andrady 2015), while others, such as fouled or high-density plastics, accumulate in high concentrations in sediments on the ocean floor (Derraik 2002; Van Cauwenberghe et al. 2013). By contrast, cod is a benthopelagic demersal fish (known as ‘groundfish’ in the Newfoundland fishing industry), meaning their main habitat is below pelagic waters but above the ocean floor (approx 150–200m below the surface) (Johansen et al. 2009). The diet of adult cod is primarily small or medium sized fish from benthopelagic waters (Cohen et al. 1990). During spawning season (early spring), cod increase their intake of benthic organisms and plant material though by late summer, which coincides with our study’s sampling season, and food intake is primarily benthopelagic fish (Cohen et al. 1990). In turn, it is possible that cod are less likely to consume plastics, given where they feed and where plastics are commonly found within the water column. Life history traits may explain the low rates of cod ingestion compared to other fish sampled in similar locations (Table 1). Further research examining the movement and location of microplastics within the water column, including the amount that is suspended from sediment into benthopelagic waters, may help to identify how and which species are most susceptible to plastic ingestion in North Atlantic waters.

Secondly, water near the coast of Newfoundland and Labrador may contain fewer microplastics than other ocean waters. Like all major bodies of water, plastics have been shown to accumulate in the North Atlantic Ocean (Eriksen et al. 2014). Newfoundland, including its coastal waters, however, is located beyond the region’s major plastic accumulation zone, the North Atlantic Subtropical gyre (Law et al. 2010). Furthermore, the waters surrounding Newfoundland receive their input from Arctic waters flowing southerly via the Labrador Current (Loder et al. 1998). While plastic ingestion has been recorded in multiple seabird species that migrate between the Arctic and North Atlantic oceans (Mallory et al. 2006; Mallory 2008; Provencher et al. 2009; Provencher et al. 2010; Provencher et al. 2014) thus indicating the presence of marine plastics within the general area of our study site, the actual plastic concentration in waters along the Newfoundland and Labrador coastline has not yet been quantified. Recent studies have suggested that Arctic sea ice may act as a sink for microplastics, as low-density plastics tend to accumulate in higher density seawater as ice freezes (Obbard et al. 2014; Lusher et al. 2015b). Microplastics are found in sea ice at six times the concentration of surrounding waters, much of which accumulates in permanent sea ice and is not released during annual sea ice events (Obbard et al. 2014). Waters flowing from the Arctic may therefore contain less microplastics than other ocean waters because of these additional sinks. This may be particularly relevant to the Newfoundland cod fishery, as larger cod are often associated with colder temperatures (Cohen et al. 1990; Björnsson and Steinarsson 2002)—something that was confirmed in the Newfoundland context through observations by local fishermen. Furthermore, the Labrador current receives inputs from Greenlandic glaciers (with icebergs regularly appearing in the spring and summer months along the Newfoundland coast) that, upon melting, further dilute microplastic concentrations with unpolluted freshwater. A similar phenomenon was suggested by Lusher et al. (2015b), who found lower than expected concentrations of microplastics in the Barents Sea, possibly due to freshwater inputs of Arctic water. However, we caution that this finding may not apply to all Newfoundland waters where cod is caught as part of the commercial and recreational cod fishery. Compared to northerly and north-easterly waters where this study took place, waters along the southern coast of Newfoundland may have higher concentrations of plastic pollution, as they receive additional inputs from the Gulf of St. Lawrence (Han et al. 1999) and wind may push pollutants from the Gulf Stream into southern waters. Future research should examine plastic ingestion in this southern area.

Third, Canada’s Northern Atlantic and Arctic regions have small and dispersed populations that contribute to relatively less onshore litter compared to other sites where cod has been sampled. Newfoundland and Labrador has a relatively low population of 530,000 people spread over 400,000km^2^, which may account for some of the difference between our results and Foekema et al. (2013) who examined cod in the much more populated North Sea. In their case, of 67 cod caught near coastal waters, 14.9% had ingested plastics, whereas none of their 13 cod caught offshore had ingested plastics—a finding that is similar to that of the present study where 0% of 17 offshore cod contained plastics. Foekema et al. (2013) attributed the greater number of plastics found near inshore waters to higher levels of local plastic pollution due to coastal proximity. Similar results were obtained by Bråte et al. (2016) in Norwegian waters, where their 302 Atlantic cod from six sampling sites around the country had a 3% ingestion rate, but cod from the Bergen City Harbour, their most populated test site, had a 27% ingestion rate.

While it is difficult to trace plastics to the original point source of pollution, we are confident that most of the plastics in the present study (n= 3, 60%) originate from the province of Newfoundland as evidenced by low levels of erosion and lack of discolouring. Newfoundland is a remote province, and plastics traveling from afar would have to endure significant time at sea. Plastics that have spent significant time at sea or on beaches show signs of wear, erosion, or fouling (Corcoran et al. 2009), which were absent from these plastics. The two plastic threads found in cod (see Image 1) are the same type and colour of plastic that is commonly found on local shorelines and coastal sea floors (Claereboudt 2004; Zhou et al. 2011), and is associated with the bottom trawls used in the area. Local waste and sewage management practices, geographical features, and proximity to the main urban area of the province likely impacts the type and incidences of plastics found in our study. Most of this study’s cod were caught at Petty Harbour and Portugal Cove, which have relatively large and open bays compared to other sampling locations, such as Belle Island and Quidi Vidi—both of which are located in small inshore bays and whose cod were found to contain plastics. Further sampling of cod in these and other inshore locations might yield larger plastic incidence rates.

A potential issue associated with comparing plastic ingestion rates among studies is the use of different lower detection limits. As has been discussed, the lower detection limit used in this study was 1mm for ‘large microplastics’, as this has been shown to be the largest size that can reliably detected visually through a compound microscope, and is the limit most commonly used in plastic ingestion studies (Song et al. 2015). However, other detection limits have been used in fish ingestion papers, where plastics of less than 1mm are included in study results (for example, Lusher et al. 2013; Phillips and Bonne 2015). This in turn might lead to an overestimation of plastics in certain species, where a greater number of small and medium sized microplastics are detected, or underestimation of others, including the present study where smaller microplastics are not investigated. We found no fibers, for example, while in studies such as Rochman et al. (2015) fibres were plentiful. This may have implications, where researchers are assessing what species are most vulnerable to plastic ingestion, and indicates the importance of using comparable methods in both accredited and citizen science. The development of reliable citizen science methods is particularly important in Newfoundland and Labrador due to the importance of fish in the local economy, community concerns regarding plastic pollution and the wellbeing of marine life, low research infrastructure, and a wealth of local knowledge by fishermen in the area. The present study contributes to ongoing efforts by marine plastic researchers to use and standardize citizen science methods.

Other studies in marine plastic pollution have used volunteer participation of citizens (citizen scientists) to contribute information, data, and samples to scientific studies (Tudor & Williams 2001, Bonney et al. 2009, Bravo et al. 2009, Ogata et al. 2009, Ribic et al. 2010, Hidalgo-Ruz & Thiel 2013, Eastman et al. 2014, Smith & Edgar 2014, Hidalgo-Ruz & Thiel 2015). In an overview of citizen science projects involving marine debris, Hidalgo-Ruz and Thiel (2015) found that 68% of studies examined the spatial distribution and composition of marine litter through beach clean ups and shoreline studies. They found that only 18% of citizen science studies addressed interaction of plastics with biota, and only one dealt with ingestion of plastics by fish, where plastics were gathered from shorelines and appraised for bite marks (Carson 2013). To our knowledge, ours is the first study that uses citizen scientists to gather GI tracts for biomonitoring and analysis. Citizen scientists, and fish harvesters in particular, are an important population for collaboration in ingestion studies because their participation allows us to sample human food webs directly. All fish in our sample were eaten by humans. Also, compared to studies where whole fish are bought from market and analyzed (such as Rochman et al. 2015a), we are able to obtain more fish with additional data, including where they were caught and under what conditions (hand line versus trawl, and nearshore versus offshore, for example). Finally, in addition to gains in sampling, citizen science also allows for input from the community. We hosted a public meeting of our results in Petty Harbour, where many of our samples were collected, before submission for publication. Citizen scientists and members of the public gave us feedback as to whether our results aligned with their own understandings of plastics and fish in the area, and they advised that we look at mackerel and capelin, two pelagic fish species also consumed in the area. They also invited us back for the following year to continue our study. For remote areas with large coastlines, in fishing communities, and on topics of public concern such as marine plastics, citizen science is an ideal methodology.

Based on this study, we will continue to monitor the food fishery in Newfoundland’s east shore in coming years to establish a more robust monitoring system, and we will look to sample the south shore of the island where plastics from the Gulf Stream are likely to occur. We will also add additional protocols to our citizen science collection to ensure entire GI tracts, rather than just stomachs, are gathered in future studies.

## Contributions

Max Liboiron is the principle investigator for this project and coordinated the study, wrote field and laboratory protocols, trained students and participated in all field and laboratory protocols, analyzed plastic samples, wrote, edited and revised the final article, and facilitated the public meeting at Petty Harbour. Liboiron is the PI on all grants that supported this study.

France Liboiron conducted the majority of laboratory wet lab work, including the analysis of plastics, wrote, edited, and revised the final article, and attended the public meeting at Petty Harbour.

Emily Wells conducted wet laboratory work, including the analysis of plastics, prepared the majority of the literature review on ingestion rates, wrote, edited, and revised the final article, and attended the public meeting at Petty Harbour.

Natalie Richárd conducted field collection and laboratory wet lab work, including the analysis of plastics, wrote and edited the final article, and attended the public meeting at Petty Harbour.

Alex Zahara wrote, edited, and revised the final article, with an emphasis on the discussion, and attended the public meeting at Petty Harbour.

Charles Mather conducted field collection, analyzed plastics in the laboratory, edited the final article, and co-facilitated the public meeting at Petty Harbour.

Hillary Bradshaw conducted field collection and attended the public meeting at Petty Harbour. Judyannet Murichi conducted field collection and attended the public meeting at Petty Harbour.

## Acknowledgements

This research was made possible through a Social Science and Humanities Research Council (SSHRC) Insight Development Grant (#430-2015-00413) and Marine Environmental Observation Prediction and Response Network (MEOPAR) grant, and the generosity of Dr. Yolanda Wiersma, Department of Biology, Memorial University of Newfoundland for dedicated laboratory space. Alicia Poole and Louis Charron provided assistance in the field and for fact-checking. We wish a special acknowledgement to all citizen participants who helped us gather samples, especially Emily Pretty, who attends Frank Roberts Junior High School in Conception Bay South and spent many hours helping us collect data. Finally, thank you to the anonymous reviewer whose comments strengthened this paper.

